# Direct from the Seed: An Atomic-Resolution Protein Structure by Ab Initio MicroED

**DOI:** 10.1101/2025.07.03.663097

**Authors:** Purna Chandra Rao Vasireddy, Timothy Low-Beer, Katherine A. Spoth, Devrim Acehan, Matthew R. Crawley, Michael W. Martynowycz

## Abstract

While purifying the seed protein crambin, we discovered that needles of pure protein nanocrystals formed spontaneously during the drying of a simple ethanolic purification drop. Contrary to traditional crystallography, these needles diffracted poorly using X-rays yet proved to be exceptionally well-suited for microcrystal electron diffraction (MicroED). By merging data from 58 such nanocrystals, we obtained diffraction to 0.85 Å resolution with an overall correlation coefficient of over 99% and solved the structure *ab initio* using a five-residue helical fragment to initiate density modification. The resulting map was of exceptional quality, enabling fully automated model building and resolving individual hydrogen atoms. This work represents the highest-resolution protein structure (0.85 Å) determined from spontaneously formed protein nanocrystals and is the first *ab initio* structure of crambin solved by electron diffraction. Our workflow demonstrates that complex biological matrices can be mined directly for sub-ångström protein structures, establishing a practical and scalable pipeline from raw biomass to atomic-level models of previously intractable targets.

## Introduction

Crystallography, the cornerstone method for determining protein structures, has traditionally been limited by crystal size. X-ray crystallography, the traditional gold standard, demands relatively large, well-ordered crystals - typically tens to hundreds of micrometers across. Yet many proteins naturally form only extremely tiny crystals, far smaller than this threshold, leaving a significant fraction of important biological molecules structurally inaccessible. Single-particle cryo-electron microscopy (cryo-EM), another powerful approach, struggles to achieve high-resolution structures of proteins less than 100 kDa.^1^ This limits its utility for many biologically important molecules, e.g. the median size of a human protein is only about 40 kDa.^2^

Microcrystal electron diffraction (MicroED) has emerged as a transformative approach to overcome these limitations.^3^ Instead of X-rays, MicroED uses electrons, which interact far more strongly with matter.^4^ This powerful interaction allows high-resolution structure determination from crystals thousands of times smaller by volume than those required for traditional X-ray methods. Crystals invisible to the naked eye,^5^ and often too delicate or too small for standard crystallography, can now yield atomic-resolution structures. This opens the door to streamlined ‘powder-to-structure’ workflows, where complex sample optimization is replaced by direct analysis of nanocrystalline material.^6,7^

Despite its promise, phasing novel protein structures by MicroED remains heavily reliant on pre-existing knowledge. Molecular replacement (MR), first demonstrated with lysozyme in 2013-2014,^8,9^ is the workhorse method and has delivered important structures like R2lox,^10^ MyD88,^11^ and the human lens membrane protein MP20.^12^ The arrival of AlphaFold2^13^ has further expanded the reach of MR. However, this reliance on homologous models creates a fundamental limitation: it generates phase bias and leaves proteins with novel folds entirely inaccessible. Establishing reliable ab initio phasing methods that remove this dependence is essential. While this has been routine for small-molecule electron crystallography for years,^6,14–16^ progress for proteins has been incremental, moving from direct methods on small peptides,^17^ to fragment-based searches,^18^ and recently to density-modification-based bootstrapping from minimal ideal fragments.^19^ Our work confronts this challenge directly, providing a robust, atomic-resolution benchmark for this latter approach, of which there are still very few examples.

In this study, we bridge this methodological gap by using the classic benchmark protein crambin.^20^ For decades, crambin has served as a proving ground for the most advanced crystallographic methods. Its 1981 structure by Hendrickson and Teeter was a landmark in *de novo* phasing using sulfur anomalous scattering,^21^ and it was later the first protein solved by Hauptman and Weeks’ purely mathematical direct methods, a triumph developed at our home institutions. In subsequent years, synchrotron and neutron^22^ sources pushed its resolution to an extraordinary 0.48 and 0.54 Å,^23,24^ making it a key model for studying ultra-high-resolution features.

Despite this storied history in X-ray and neutron science, a complete *de novo* investigation of crambin by MicroED has remained unexplored. While a 1.07 Å MicroED structure solved by molecular replacement has been briefly described in a technical note (Rigaku Application note PX031),^25^ the full potential of using electrons to solve this benchmark protein from first principles has not been realized. We discovered that a simple ethanolic extraction from ground seeds spontaneously yields an abundant supply of nanocrystals ideal for such a test. Therefore, we set an ambitious goal: to determine if data from these naturally formed nanocrystals were of sufficient quality to not only break the sub-ångström barrier but to solve the structure entirely *ab initio* from a minimal, non-homologous five-residue helical fragment.^26^

Successful *ab initio* structure determination would represent the first electron-based atomic-resolution structure of this pivotal protein, validating a streamlined pipeline from raw biological material directly to detailed structural models. This result not only provides an essential benchmark for advancing electron crystallography but also demonstrates a practical, scalable approach capable of unlocking structural information from previously underexplored biological targets, such as naturally occurring macromolecular products.

## Results and Discussion

### A Tale of Two Crystals: Swapping Fortunes in X-ray and Electron Diffraction

Our investigation began with a serendipitous discovery. During a simple ethanolic extraction of crushed *Crambe abyssinica* seeds (**Supplementary Figure 1**), we found that a 1 µL droplet of the protein-rich solution nucleated a dense shower of sub-micron, needle-shaped crystals within seconds as the solvent evaporated (**Figure 1A**). No crystallization screen is required. The process is rapid and spontaneous. This shortens the entire pipeline from raw seed to refined atomic model. The process can be completed in a week or less. These needles proved unsuitable for X-ray diffraction. A single 14 µm needle diffracted poorly, only reaching 1.6 Å resolution and suffered from rapid radiation decay (**Supplementary Figure 2, Supplementary Table 1**). In contrast, they proved ideal for MicroED. When loaded into the TEM, we immediately noticed many tiny rod-shaped crystals on the grid (**Figure 1B**). High resolution imaging of these crystals showed clear lattice lines (**Figure 1C**) that contained information to 3.6 Å by analysis of the Fourier transform of these images (**Figure 1D**). These crystals consistently diffracted electrons to high resolution (**Figure 1E**).

**Figure 1.**
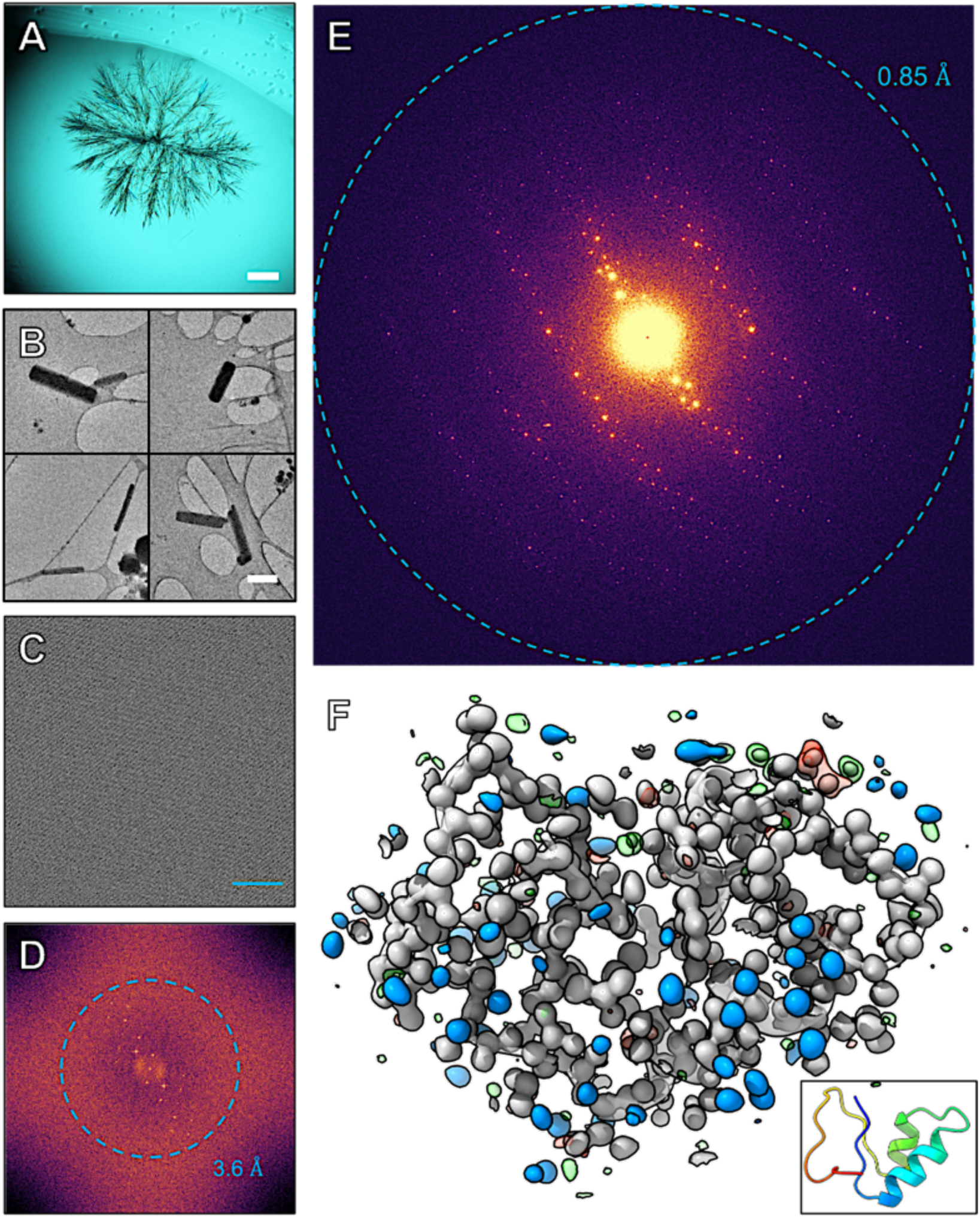
Characterization of crambin microcrystals. (**A**) Crambin protein crystals crashed out in a drop on a cover slide. Scale bar 125 μm. (**B**) Collection of four TEM images at 4300 × magnification showing small rod-shaped crystals of crambin. Scale bar 1 μm. (**C**) High resolution TEM image of the crystal lattice of a typical crambin crystal in the TEM. Scale bar 200 Å. (**D**) Fourier transform of panel C showing lattice order out to 3.6 Å. (**E**) 1° of MicroED data showing diffraction to 0.85 Å resolution. (**F**) The final refined model of crambin as a transparent cartoon. The 2Fo-Fc map is in gray and contoured to 1.5 σ. The blue densities are the same map segmented around the modelled water residues. The Fo-Fc map with a threshold at +3 σ is in green and with a threshold of −3 σ is shown in red. A cartoon depiction of the final model is inset in the same orientation.

We also grew large, block-like crystals using traditional vapor diffusion (**Supplementary Figure 3**). These crystals exhibited the opposite behavior: they produced excellent X-ray diffraction to beyond 0.8 Å but were initially considered a secondary choice for MicroED due to their thickness. Initially, we mechanically crushed these high-quality blocks to obtain fragments small enough for MicroED. The rationale was twofold: first, we hoped to leverage their superior internal order observed with X-rays; second, we anticipated that randomly oriented fragments would mitigate the severe preferred orientation expected from the needle morphology, thereby yielding a more complete dataset. However, this approach failed. The crushed fragments produced markedly inferior electron diffraction data compared to the spontaneously formed needles (**Supplementary Figure 4**).

This surprising inversion of crystal quality between the two diffraction methods became a central finding of our work. The gentle, in-solution self-assembly of the nanocrystals preserves a higher degree of internal order for electrons, while the block-like crystals that excel with X-rays are damaged by the mechanical stress of fragmentation. This led us to a critical decision: instead of pursuing randomly oriented but damaged crystal fragments, we would tackle the preferred orientation problem of the spontaneously formed needles directly using a serial crystallography approach. By merging data from dozens of crystals, we aimed to overcome geometric data loss inherent to a single orientation and build a complete, high-redundancy dataset.

### Achieving an Atomic-Resolution MicroED Dataset by Overcoming Anisotropy

Having established the superiority of the spontaneously formed nanocrystals for MicroED experiments, we proceeded with data collection. The primary technical hurdle was, as anticipated, the severe preferred orientation of the needles on the grid.^27^ This morphology introduces a fundamental challenge for MicroED: diffraction quality is highest at low tilt angles where the electron beam passes through the thinnest dimension of the crystal but degrades at high tilts as the effective crystal thickness increases,^28^ compounding multiple scattering and absorption effects. This inherent tilt-dependent resolution decay is a major source of anisotropy in data from plate- or needle-like crystals.^10^

To obtain a high-quality dataset, we merged data from 58 individual nanocrystals using a serial MicroED strategy. To maximize data completeness and mitigate the “missing wedge” effect inherent to single-crystal collection, we collected a 30° or 60° data wedge from each crystal. The starting angle for each wedge was intentionally varied, ensuring that data from different crystals sampled unique regions of the full goniometer tilt range. Even with this comprehensive collection strategy, the merged dataset exhibited the expected anisotropy. A standard spherical resolution cutoff would therefore be suboptimal, either conservatively low, discarding valuable high-resolution data, or too high, including noise from the poorly ordered directions.

To produce the most physically meaningful and high-quality dataset, we chose to explicitly address this anisotropy. We processed the integrated intensities with the STARANISO server,^29^ which determines the direction-dependent resolution limits and applies a corresponding elliptical truncation. This process correctly removes the weak, high-resolution impaired reflections from the poorly diffracting directions while preserving the strong, high-resolution data from the well-ordered directions. While modern phasing programs like Phaser are robust against some noise, providing them with the cleanest possible dataset maximizes the chances of success in a challenging *ab initio* case. Similarly, refinement benefits from a dataset where resolution limits are physically justified.

The final, anisotropy-corrected dataset extended to an overall resolution of 0.85 Å and exhibited high internal consistency and an effective (ellipsoidal) completeness of 98.6% (**Table 1**). The high data multiplicity of 42.67, achieved by merging data from 58 nanocrystals, was a critical factor in producing the exceptional signal-to-noise ratio within this well-defined resolution ellipsoid. The diffraction limit is defined by an elliptical cutoff with resolutions of 0.84 Å, 0.85 Å, and 1.49 Å. As recent work by Xu et al. has shown,^30^ high multiplicity can significantly improve the overall quality of merged MicroED datasets, and our result provides a powerful validation of this strategy. Analysis of the data showed no evidence of specific radiation damage.^31^ No significant difference map peaks appeared at susceptible sites such as disulfide bonds or acidic residues.

**Table 1.** MicroED structure of crambin.

| **Data collection** |  |
| --- | --- |
| No. of crystals | 58 |
| Space group | *P* 2<sub>1</sub> |
| Cell Dimensions<br>(a, b, c) (Å)<br>(α, β, γ) (°) | 41.490, 18.790, 22.520<br>90.00, 90.84, 90.00 |
| Resolution (Å)<sup>1</sup> | 22.58 – 0.85 (0.87 - 0.85) |
| R<sub>pim</sub> (%) | 3.9 (48.2) |
| < I / σ I > | 12.118 (1.00) |
| Completeness (%) | 98.6 (83.6)<sup>1</sup> |
| Redundancy (#) | 42.67 (8.60) |
| **Refinement** |  |
| Resolution (Å) | 22.58 – 0.85 (0.872 -0.85) |
| No. Unique Reflections (#) | 22,022 (619) |
| R<sub>work</sub> / R<sub>free</sub> (%) | 15.4 / 17.2 |
| No. Atoms / B-factors | 426 / 6.68 |
| Protein (# / Å<sup>2</sup>) | 354 / 4.60 |
| Water (# / Å<sup>2</sup>) | 42 / 16.89 |
| **RMS deviations** |  |
| Bond lengths (Å) | 0.021 |
| Bond Angles (°) | 1.96 |
| Clash score | 2.92 |

<sup>1</sup> Statistics after elliptical truncation and scaling with the STARANISO server.

### Ab Initio Phasing and Automated Structure Completion

The central achievement of this work was to solve the structure without a homologous model. Our workflow, conceptually similar to the fragment-based Fragon^32^ pipeline developed for X-ray data, began with an attempt at molecular replacement in PHASER^33^ using a generic five-residue poly-alanine helix. This yielded a statistically marginal solution (TFZ = 7.2) with an uninterpretable potential map (**Figure 2A**); however, these initial phases were sufficient to bootstrap the dynamic density modification algorithm in ACORN.^34^ This fragment-based phasing approach is similar to the strategies recently used to solve the structures of lysozyme and Proteinase K by MicroED,^19^ confirming its power for solving *ab initio* protein structures from high-quality electron diffraction data. The overall concept is also similar to the ARCIMBOLDO^35^ and SHELXE^36^ based approaches. This approach solved the phase ambiguity, producing an exceptionally clear map in which the entire polypeptide chain was visible (**Figure 2A**). The quality of this *ab initio* map was high enough for BUCCANEER^37^ to automatically build 100% of the crambin model in a single pass. The final structure was refined against the twinned, anisotropy-corrected data to R_work_ / R_free_ values of 15.4% / 17.2%, yielding a model with excellent stereochemistry (**Table 1**). The resulting 2F_o_-F_c_ map resolves all non-hydrogen atoms and many hydrogen atoms, confirming the genuine atomic resolution of the structure (**Figure 1F**).

**Figure 2.**
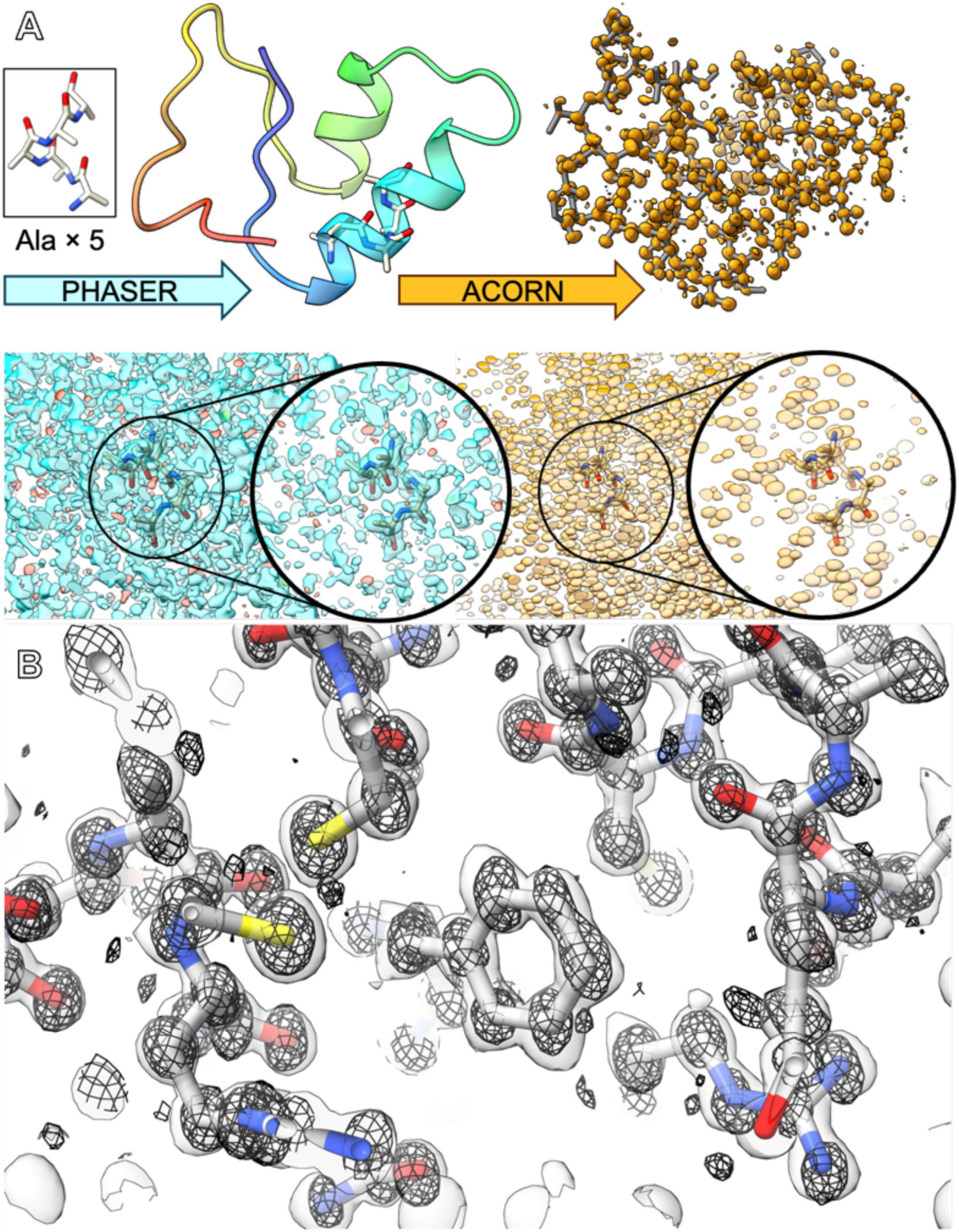
*Ab initio* structure solution of crambin at atomic resolution by MicroED. (**A**) Pipeline showing how the structure was phased. An ideal helical fragment consisting of 5 alanine residues (**top left, inset**) was placed in the unit cell by PHASER. The location of the placement relative to the final structure is shown in the **A, top left**. The resulting map could not be understood or built into outside of the placed fragment (**A, bottom, blue density**). The placed fragment was used to generate initial phases for phase improvement by the density modification program ACORN, which returns a fully interpretable map (**A, right, orange densities**). This map was used to automatically build the entire protein using BUCCANEER (**A, top right**). (**B**) The final refined model and 2Fo-Fc map (**B, surface**) are shown with the original |E| map from density modification, demonstrating the atomic positions (**B, mesh**).

Crambin extracted from seeds contains two natural isoforms known as PL and SI, differing at residues 22 and 25.^38^ Our workflow did not separate these variants; however, the exceptional quality of our final map allowed us to resolve this heterogeneity directly. Refinement of both isoforms with partial occupancies converged to a stable, near 50:50 ratio and, critically, eliminated all significant positive or negative peaks in the difference maps at the sites of variation, confirming the accuracy of the mixture model.

### Forging the Next Link in a Benchmark’s Legacy

By solving the first *ab initio* electron structure of crambin, our work forges a direct link between the foundational phasing breakthroughs of the 20^th^ century and the new frontier of atomic-resolution electron crystallography. The original solutions of crambin by Hendrickson, Teeter, Hauptman, and Weeks were not merely structural triumphs; they were powerful proofs of principle that validated new phasing methodologies for the entire field.^21,39^ This role as a methodological testbed has continued, including its use as a target for total chemical synthesis to validate protein ligation and folding strategies.^40^ We believe our result serves the same purpose for the modern era of MicroED.

We have shown that a minimal, non-homologous fragment is sufficient to phase atomic-resolution data from nanocrystals obtained without deliberate crystallization. This approach provides crystals of significantly higher quality than those reported from chemically synthesized material.^40^ This achievement transforms crambin from being solely a benchmark for X-ray^24^ and neutron^22,41^ methods into a vital, cross-platform standard. The availability of our 0.85 Å electrostatic potential map alongside the classic electron-density and neutron-scattering-length maps now provides an invaluable resource for calibrating and developing the next generation of electron-specific refinement tools, especially those designed to tackle the grand challenges of dynamical scattering^42^ and charge density analysis.^43^

We have determined an atomic-resolution electron diffraction structure of crambin, at 0.85 Å. The structure was solved *ab initio* using nanocrystals that formed spontaneously, without deliberate crystallization. Although our fragment-based approach differs from sulfur anomalous dispersion or traditional direct methods, it shares their underlying principle: starting from minimal initial information to build a complete structure.

The resulting electrostatic potential map shows precisely how electrons scatter off the Coulomb potential of every atom.^44^ Unlike X-ray maps, which highlight electron clouds, or neutron maps, which locate atomic nuclei,^45^ MicroED maps directly show the electrostatic potential felt by incoming electrons. This potential is sensitive to both the number of electrons and the atom’s net charge. As a result, map peaks for negatively charged atoms appear less intense than expected based on atomic number alone. Meanwhile, even individual hydrogen atoms appear clearly as positive peaks,^46^ rather than being invisible or merely inferred.

Since our map provides a complementary chemical picture of crambin to those available through X-ray or neutron techniques,^47^ crambin once again serves as a benchmark, this time as a gold-standard reference for future electron-specific methods, such as modeling electron scattering, analyzing charge density, and accurately visualizing hydrogen bonds.^46^ In this way, our work connects past breakthroughs in solving the crystallographic phase problem with today’s emerging field of atomic-resolution electron crystallography.

## Conclusions

In this work, we have demonstrated a complete and streamlined pipeline for the *ab initio* structure determination of a protein to atomic resolution using MicroED. We have shown that sub-micron protein needles, formed spontaneously during a simple ethanolic extraction, can be carried directly from crude plant material to a fully refined 0.85 Å structure without the need for extensive crystallization trials or a homologous search model. This low-tech crystallization, high-impact workflow potentially turns raw biological matter into ready-made nanocrystal libraries, enabling structural analysis of historically intractable proteins.

Our findings significantly lower the barrier for obtaining high-resolution structures, particularly for proteins that crystallize only as sub-micron needles or lack known structural homologues. The high redundancy of our merged dataset not only contributed to the quality of the final model, as previously suggested by others,^30^ but also points toward future data collection strategies. We believe that future experiments employing serial electron crystallography^48^ by scanning the beam or the stage with extremely short exposures could further improve data quality by eliminating stage rotation artifacts entirely and vastly increase multiplicity.

The exceptional quality of this crambin dataset provides an ideal benchmark for future electron-based refinement methods. Notably, our R_work_ / R_free_ values (15.4%/17.2%) are among the lowest reported for a protein structure by MicroED, and the data quality metrics (R_pim_ = 3.9%) are consistent with largely kinematic scattering. This is likely due to the use of extremely thin nanocrystals (<100 nm), a highly parallel electron beam, and the use of highly redundant data, which likely minimize any dynamical effects.^28^ Nevertheless, no protein structure has yet been refined with a full dynamical model. Our 0.85 Å data, with its high redundancy and resolution, represents a perfect testbed for developing and validating such tools, which will be essential for pushing the boundaries of accuracy.

The robustness of this system makes it an ideal platform for systematically studying the effects of different solvents, cryoprotectants, or ligands on protein structure at atomic resolution. Furthermore, this high-quality electron-based model, when combined with existing ultra-high-resolution X-ray and neutron data, creates a unique opportunity for joint refinement,^49^ which could provide unparalleled insight into chemical bonding and protonation states. By incorporating energy filtration,^50,51^ faster detectors,^52^ and advanced sample preparation techniques like ion-beam milling,^53,54^ the stage is now set to explore hydrogen bonding networks and charge-density distributions, extending true atomic-resolution structural biology to an ever-wider range of challenging biological systems.

## Materials and Methods

### Protein Isolation and Nanocrystal Formation

Crambin was extracted from 230 g of manually de-husked and then crushed *Crambe abyssinica* seeds following a modified protocol.^55,56^ The ground kernels were first defatted with hexanes and then subjected to three rounds of extraction with 80% acetone in water (%v/v). After evaporation of the acetone under reduced pressure, the aqueous solution was stored at 4 °C for 18 hours. This solution yielded a pale-yellow solid pellet upon centrifugation, which was redissolved in a 70% ethanol-water mixture (%v/v). While attempts to grow large single crystals by slow cooling or vapor diffusion from this solution were unsuccessful, we made a key discovery during routine sample inspection. When a 1 µL droplet of the ethanolic protein solution was allowed to evaporate on a microscope slide, it rapidly nucleated a dense shower of sub-micron, needle-shaped crystals. These spontaneously formed nanocrystals, unsuitable for X-ray diffraction, became the source material for all subsequent MicroED experiments. The full detailed extraction protocol is available in the **Supplementary Methods**.

### MicroED Grid Preparation and Data Collection

Holey-carbon copper grids (Quantifoil Cu 300 R1.2/1.3) or lacey carbon grids (Cu 200, Electron Microscopy Sciences) were glow-discharged (15 mA, 30s). The grids were transferred to either a Leica GP2 plunge freezer or Thermo-Fisher Vitrobot equilibrated to 4 °C and 95% relative humidity. A 3 µL aliquot of the crystalline slurry was applied, blotted for approximately 30s, and plunge-frozen in liquid ethane. MicroED data were recorded on a Glacios TEM (200 kV, λ = 0.0251 Å) equipped with a Falcon 4 detector. Two sets of data were collected using SerialEM:^57,58^Linear-mode datasets with wedges 0.2°/frame and 1.6 e⁻ Å⁻² total exposure. Counting-mode datasets were collected with 0.1°/frame and 0.8 e⁻ Å⁻² total exposure. We did this in the hopes the low-resolution reflections in linear mode would be less prone to coincidence loss, and the high-resolution reflections in counting mode would be more accurately recorded. The total exposure per crystal that was used for MicroED data collection was kept below 2 e⁻ Å⁻². In both schemes the starting angle of each wedge was incremented stepwise so that the combined datasets sample a stage tilt range of +/− 60°. Counting mode and linear mode datasets were collected on different crystals.

### Data processing and Refinement

Raw movie frames were converted to the miniCBF format using a custom Python pipeline that performed 2×2 binning and a two-pass, smoothed beam-center correction (**see Supplementary Methods for full details**). The corrected images were processed with XDS (BUILD 20230630) and scaled with XSCALE. The final merged dataset from 58 crystals was processed by the STARANISO server to correct for anisotropy. Phasing was performed using a combination of Phaser and ACORN, followed by automated model building in BUCCANEER. The final structure was refined using REFMAC5 with explicit modeling of a 10.0% twin fraction. Final data collection and refinement statistics are provided in **Table 1**.

## Supporting information

Supplemental Video 1: Post MR Density

Supplemental Video 2: Post DM Density

## Author contributions

P.C.R.V, T.L.B, D.A, and K.A.S prepared samples. K.A.S, P.C.R.V, T.L.B, M.R.C and M.W.M collected data. P.C.R.V, T.L.B, M.R.C and M.W.M processed data. M.W.M conceived the project and directed the work. The paper was written, and figures were made by contributions from all authors.

## Acknowledgements

Seeds of *Meyer Crambe* were grown near Carrington, North Dakota and generously provided by Mike Ostlie, from the Carrington Research Extension Center at North Dakota State University. We would like to thank Max TB Clabbers, Johan Hattne, Chris Campomizzi, Edward Snell, and Sarah Kohne for helpful discussions. We would also like to thank Gabby Budziszewski, Elizabeth Snell, Tiffany Wright, and Sarah EJ Bowman from the National Crystallization Center for use of their light microscope and assistance with several crystallization tools.

## Data and Code availability

Coordinates and structure factors for crambin determined by MicroED are deposited in the PDB and maps from this work are available in the EMDB. All raw MicroED data in MRC format and associated metadata for this work have been uploaded to Zenodo. Data conversion scripts and code used in this paper are available from the PI upon request and from our GitHub page.

## Funding Information

The Martynowycz Laboratory is supported by funds from the Community Foundation for Greater Buffalo. Rigaku XtaLAB Synergy-S, a part of the Chemistry Instrument Center (CIC) at UB, was purchased with NSF CHE-2216151. The National Crystallization Center at UB HWI is supported by the National Institutes of Health (NIH), National Institute of General Medical Sciences (NIGMS) (R24GM141256).

## Supplementary Information

**Supplementary Figure 1.**
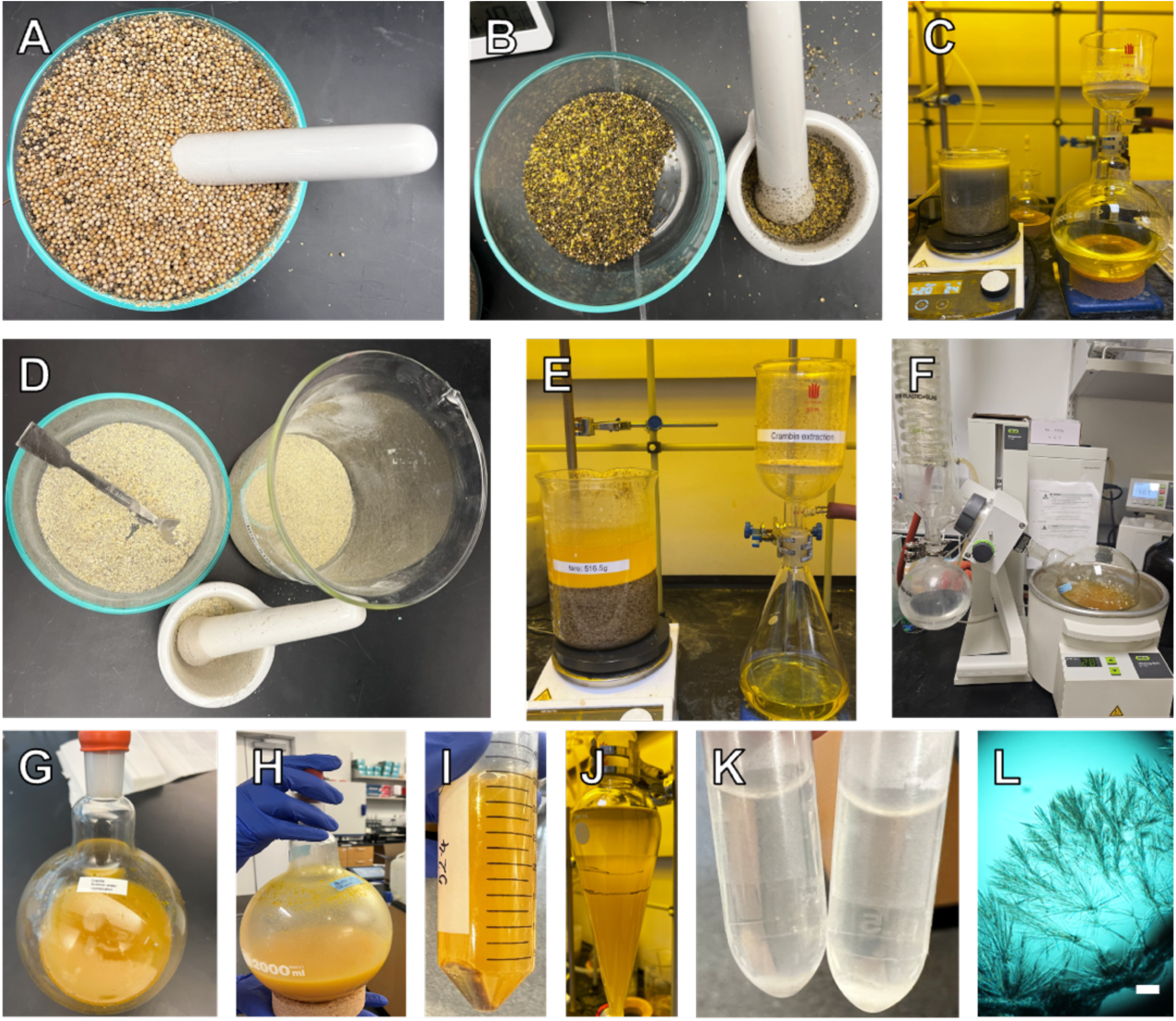
The Crambin Extraction Process. (**A**) Husking crambin seeds. (**B**) Ground seeds prior to defatting. (**C**) Defatting by mixing with hexanes and filtering. (**D**) Ground seed meal prior to crambin extraction. (**E**) Crambin extraction using 80% acetone-water by mixing and filtering. (**F**) Acetone removal under reduced pressure and <40°C. (**G**) Solution after acetone removal. (**H**) solution after 18h at 4°C, (**I**) centrifuged solution. (**J**) hexane washings to crude crambin pellet dissolved in 70% EtOH-water. (**K**) crystals obtained by slow evaporation. (**L**) microcrystals seen under light microscope. Scale bar 125 μm.

**Supplementary Figure 2.**
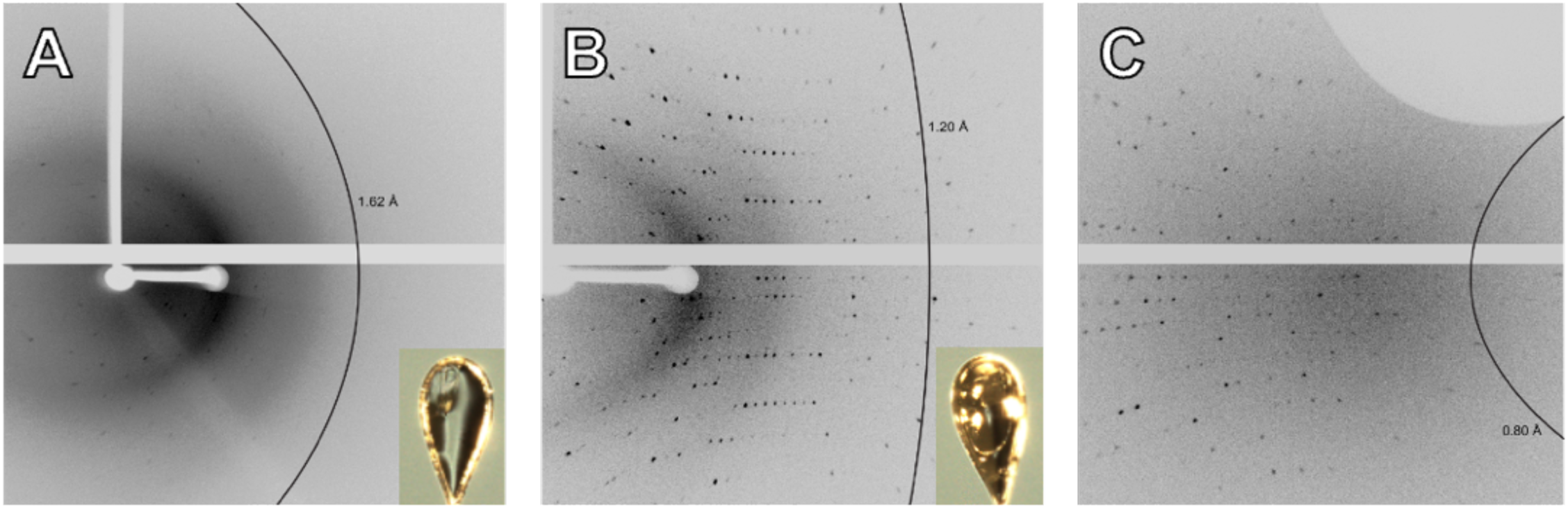
Comparison of needle shaped crystals grown by evaporation to block shaped crystals grown by vapor diffusion by single crystal X-ray crystallography. (**A**) A single frame (0.5° width) of a needle-like crystal of crambin (see inset) with a 120 second exposure time and resolution ring at 1.62 Å. (**B**) A single frame (0.5° width) of a block-like crystal of crambin (see inset) with a 10 second exposure time and resolution ring at 1.20 Å. (**C**) The same crystal from panel B with a (0.5° width) with a 35 second exposure time showing diffraction beyond the resolution ring at 0.80 Å.

**Supplementary Figure 3.**
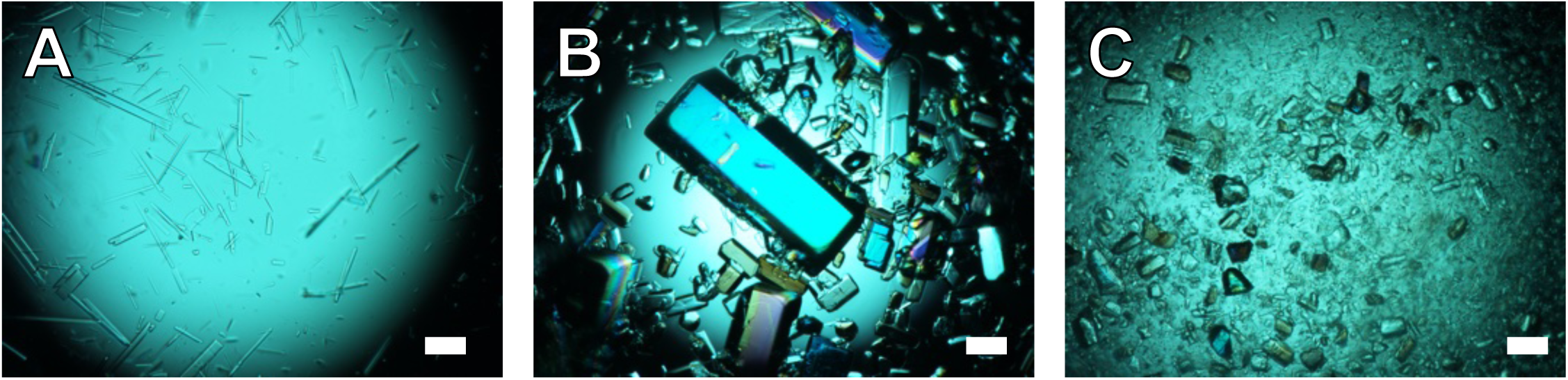
(**A**) Crashed out needles of crambin formed by slow evaporation. (**B**) large block shaped crystals of crambin with sizes up to 1 mm across. (**C**) Crystals from panel B after being mechanically crushed. All scale bars 125 μm.

**Supplementary Figure 4.**
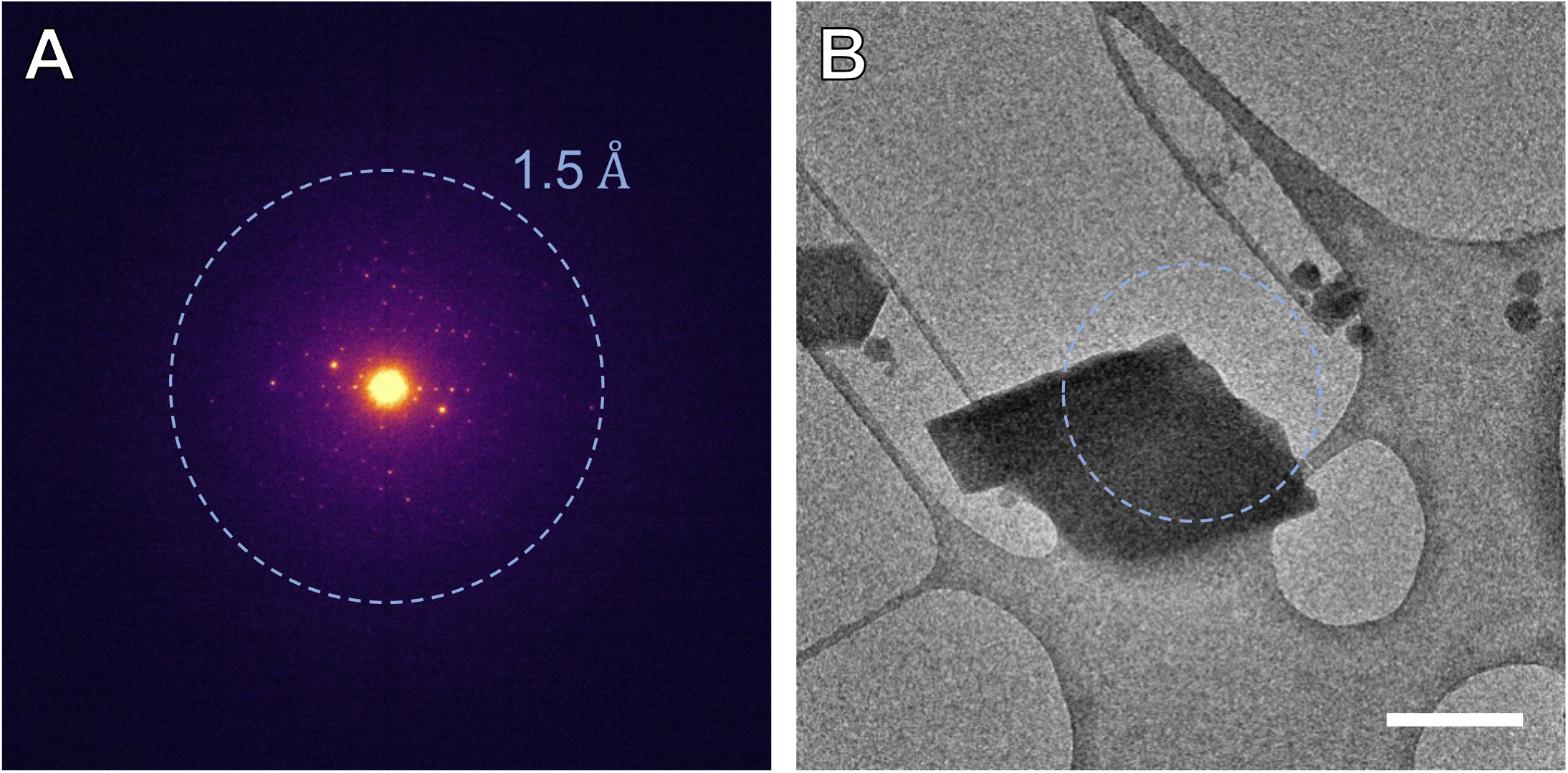
Electron diffraction from fragmented block-like crystal. (**A**) Representative electron diffraction pattern (1° width) of a fragmented block-like crystal of crambin, resolution ring at 1.5 Å. (**B**) TEM image of the corresponding crystal. Dashed circle indicates beam position and diameter during diffraction. Scale bar 1 μm.

**Supplementary Table 1.** X-ray Crystallographic details for crystals of crambin.

|  | **Plate/needle crystal** | **Block crystal** |
| --- | --- | --- |
| Temperature (K) | 110 | 110 |
| Crystal system | Monoclinic | Monoclinic |
| Space Group | *P2<sub>1</sub>* | *P2<sub>1</sub>* |
| a (Å) | 22.296(4) | 22.3196(2) |
| b (Å) | 18.476(5) | 18.4590(1) |
| c (Å) | 40.84(2) | 40.7640(4) |
| β (°) | 90.64(3) | 90.5399(8) |
| Crystal size (mm x mm x mm) | 0.211 × 0.073 × 0.014 | 0.189 × 0.135 × 0.070 |
| Radiation | Cu Kα (λ = 1.54184) | Cu Kα (λ = 1.54184) |

**Supplementary Table 2.**
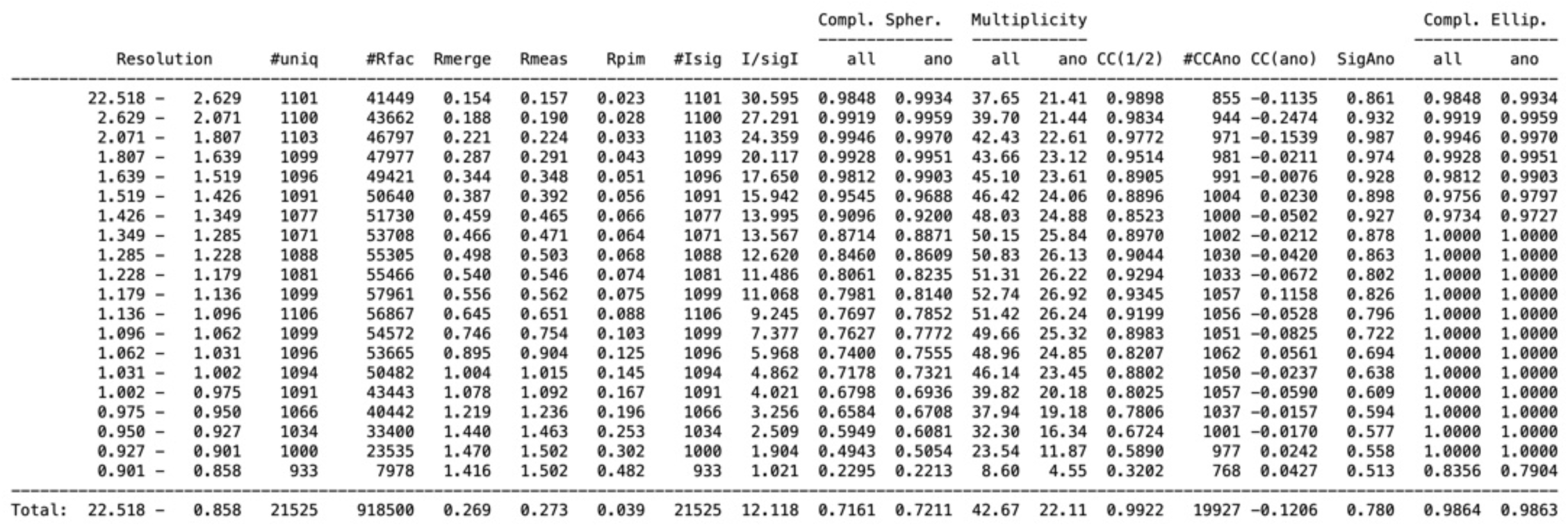
Scaling and merging statistics after the STARANISO server.

## 7. Supplementary Methods

### a) Materials

Seeds from *Crambe abyssinica* (Meyer *Crambe*, grown near Carrington, North Dakota, 2022) were provided by the NDSU Carrington Research Extension Center. Hexanes (BDH Chemicals, >98.5%), acetone (BDH Chemicals, >99.5%), and ethanol (Pharmco, 95%, 190 proof) were used as purchased. Water was purified to 18.2 MΩ·cm using a Milli-Q system. Lacey carbon (Cu 200 mesh, Electron Microscopy Sciences) and holey carbon (Quantifoil R1.2/1.3, Cu 300 mesh) grids were used without further modifications.

### b) Protein isolation and crystallization

The extraction protocol was adapted from classic literature.^55,56^ A visual summary of the steps is provided in **Supplementary Figure 1.**

*Seed Preparation:* 230 g of seeds were manually de-husked to yield 160 g of kernels, which were ground using a mortar and pestle.

*Defatting:* The powder was defatted by stirring in hexanes (4 × 750 mL) for one hour per wash, with filtration after each step. The seed meal was then air-dried for 18 hours.

*Acetone Extraction:* The dried powder was stirred in 80% (v/v) aqueous acetone (10 mL per gram of initial seed meal) for 60 minutes and filtered. This extraction was repeated twice more.

*Initial Concentration:* The combined acetone filtrates were concentrated under reduced pressure (rotary evaporator, water bath at 27-30 °C) until the solution became hazy and acetone distillation ceased. The solution was stored at 4°C for 18 hours; no precipitation was observed.

*Pellet Isolation:* The hazy solution was centrifuged (4°C, 10,000 RPM / 16,000g, for 1hr) to yield a pellet containing a pale-yellow solid and a brown gummy layer.

*Ethanolic Dissolution:* The supernatant was decanted, and the pellet was fully redissolved in ∼150 mL of 70% (v/v) aqueous ethanol. This solution was washed with hexanes (3 × 100 mL) to remove residual oils, with 5-8 mL of fresh ethanol added after each wash to maintain protein solubility.

*Spontaneous Nanocrystal Formation:* The final ethanolic solution was concentrated by half under reduced pressure (28–30 °C). While storage at 4°C did not produce crystals, we observed that when a 1 µL aliquot was placed on a glass slide, a dense field of needle-like microcrystals formed within seconds upon solvent evaporation. This crystalline slurry was used directly for MicroED grid preparation.

### c) X-ray crystallography control experiments

To benchmark the quality of the spontaneously formed nanocrystals, large single crystals of two different morphologies (plates and blocks) were grown by vapor diffusion. An aliquot of the 70% ethanol solution of crambin was passed through a neutral alumina plug to remove color and then diffused against a 50% ethanol-water solution for two weeks.

Diffracted intensities were measured on a Rigaku XtaLAB Synergy-S diffractometer with a HyPix-6000HE detector and a Cu-target PhotonJet-S microfocus source (λ = 1.54184 Å). Crystals were cryo-protected in a 1:1 mixture of mother liquor and glycerol and maintained at 110 K in a nitrogen cryostream. Data were collected, indexed, integrated, and scaled using CrysAlisPro.^59^ A single 14 µm needle, representative of the morphology used for MicroED, was tested with X-rays. It diffracted poorly to only 1.6 Å and showed signs of rapid radiation damage, rendering it unsuitable for X-ray structure determination. These were the crystals subsequently crushed for the comparative MicroED experiments. In contrast, the block-like crystals grown by vapor diffusion and later crushed for MicroED experiments diffracted to beyond 0.8 Å resolution on this X-ray diffractometer. The unit cell parameters are reported in **Supplementary Table 1**.

### d) MicroED grid preparation

Holey-carbon (Quantifoil Cu 300 R1.2/1.3) or lacey carbon (Cu 200, EMS) grids were glow-discharged at 15 mA for 30s using a Pelco Easi-Glow. Grids were transferred to either a Leica GP2 or a Thermo Fisher Vitrobot Mark IV plunge freezer. The GP2 was equilibrated to 4 °C and 95% relative humidity and the Vitrobot was set to 22°C and 100% relative humidity for 15-20 minutes prior to use. After allowing the grid to equilibrate in the chamber for 30s, a 3 µL aliquot of the crystalline slurry was applied. Grids were blotted manually from the back side for approximately 10s inside of the Vitrobot and for 30s from behind automatically in the GP2 before being plunge-frozen in liquid ethane.

### e) TEM lattice imaging

High resolution TEM images of crystal lattices were collected at a nominal magnification of 120kx corresponding to a pixel size of 0.851 Å using a Thermo Fisher Glacios operating at 200kV at liquid nitrogen temperature. A Falcon 4 direct detector operating in counting mode was used to save the data in EER format with an exposure time of 3.48s corresponding to 812 EER frames. The total exposure was ∼37 e⁻/Å². MotionCor2^60^ version 1.6.4 was used to align the movie frames using 116 fractions, 1x EER upsampling, and no dose weighting applied.

### f) MicroED data collection

MicroED data were recorded on a Thermo Fisher Glacios transmission electron microscope operated at 200 kV (λ = 0.0251 Å) at liquid nitrogen temperatures. A Falcon 4 direct electron detector was used for data acquisition under the control of SerialEM.^57,58^ The microscope was set to microprobe mode with a 20 µm C2 aperture and no SA aperture, resulting in a parallel beam of approximately 2 µm diameter. A two-part strategy was employed to maximize the dynamic range: half of the datasets were collected in linear mode and half in counting mode. For linear-mode wedges, the total exposure was ∼1.6 e⁻/Å² over a 60s collection with continuous stage rotation, resulting in a tilt of 0.20° per 1-second frame. For counting-mode wedges, the total exposure was ∼0.8 e⁻/Å², with a finer slicing of 0.10° per 1-second frame. The total accumulated dose per crystal was kept below 2.0 e⁻/Å². Starting angles of −60°, −30°, and 0° were used for collection of 60° linear-mode wedges. Counting-mode 30° wedges were collected with starting angle varying from −60° to +30° in 15° increments.

### g) Conversion of MRC movies to miniCBF format

Raw movie stacks in MRC format were converted to the miniCBF format suitable for crystallographic processing using a custom Python script, which leverages the FabIO,^61^ NumPy,^62^ and SciPy^63^ libraries. This pipeline performed several critical pre-processing steps.

*Metadata Extraction:* For each movie, experimental parameters were parsed from the corresponding .mdoc file. These included the unbinned camera pixel size (14.0 µm), accelerating voltage (200 kV), camera length (961.06 mm), and frame-by-frame goniometer rotation angles.

*Binning:* The raw 4096 x 4096 (or 2048 × 2048 if hardware binned) pixel frames were binned by a factor of 4 (or 2) in both X and Y dimensions, producing final 1024 × 1024 pixel images with an effective pixel size of 56 µm.

*Pedestal Correction:* To ensure all pixel counts were positive in the linear mode datasets, a pedestal value was automatically calculated by finding the minimum integer value in the entire image stack (e.g., −159 for one dataset) and adding its absolute value to every pixel.

*Beam Center Correction:* A two-pass algorithm was applied to correct for beam drift during data collection. In the First Pass / Peak Finding step, an initial beam center was located by applying a Gaussian blur (σ=3.0) to a 100×100 pixel central region of interest and fitting a 2D Gaussian function to the resulting peak for each frame in the movie. The second pass consists of Smoothing and Validation. Any center that jumped more than 2.0 pixels from its predecessor was marked as an outlier. A Savitzky-Golay smoothing filter (window length=11, polynomial order=2) was then applied to the path of the valid centers. Outliers and missing centers were filled in using the smoothed path. Finally, each frame was shifted to a common, smoothed center using linear interpolation before being written to a miniCBF file with a complete header.

### h) Data integration, scaling, and anisotropy correction

Corrected CBF files were processed with XDS^64^ (BUILD 20230630). Datasets were initially indexed in space group *P* 1, *P* 2 or *P* 2 2 2. The resulting INTEGRATE.HKL files were analyzed with POINTLESS (CCP4 9.0) to determine the correct lattice symmetry, which was consistently identified as monoclinic, space group P2₁. Each dataset was then re-integrated in XDS using the correct symmetry and a fixed reference unit cell.

From an initial pool of over 77 crystals that showed diffraction, a final selection of 58 datasets that showed high mutual correlation coefficients and consistent cell parameters were scaled together using XSCALE^65^ after clustering and pruning down using XSCALE_ISOCLUSTER (KD 2024-10-27). The final scaling process used a target unit cell of a = 41.49 Å, b = 18.79 Å, c = 22.52 Å, β = 90.84°-the median of the cell values used in the scaling and merging steps. The resulting merged reflection file, XSCALE.1.HKL, contained 937,603 total observations of 26,147 unique reflections and was used as the input for anisotropy analysis using the STARANISO^29^ server. The full scaling and merging statistics for the final dataset are reported in **Supplementary Table 2.**

### i) Phasing, Model Building, and Refinement

*Ab initio Phasing and Automated Building:* Phasing was performed using a multi-step procedure. First, an ideal 5-residue α-helix (theor-helix-5.pdb from the CCP4^66^ fragment library) was placed using molecular replacement in PHASER^33^ (via the Phenix command-line utility). The best solution yielded an LLG of 64 and a TFZ of 7.2. The phases from this solution were used to initiate density modification in ACORN.^34^ ACORN was run with default parameters, terminating after 48 cycles and reaching a map correlation of 68.9%. The resulting map was used as input for BUCCANEER,^37^ which built the complete model in one pass. Based on the clarity of the density modified and refined density maps in regions of known microheterogeneity (residues 22 and 25),^38^ the model was built and refined using both potential side chains which appeared clearly in the difference densities and the occupancies were allowed to refine, resulting in an approximately 50/50 mixture of the PL and SI isoforms.

*Crystallographic Refinement:* The final model was refined in REFMAC5^67,68^ (v5.8.0430) using electron scattering factors. The protocol involved 5 cycles of manual model adjustment in Coot followed by automated refinement. Psuedo-merohedral twinning^69^ was modeled using the TWIN -h -k l instruction with a refined twin fraction of 10.0%. Anisotropic B-factors and riding hydrogen atoms were included in the final rounds. Final model statistics and validation metrics from MolProbity^70^ are reported in **Table 1**.

### j) Figure generation

Plots were generated using Python 3.11. Diffraction patterns and sums were produced using FIJI.^71^ Figures were arranged using PowerPoint. Density figures and models were prepared using UCSF ChimeraX^72^ 1.9 and PyMol^73^ 3.1.

**Supplementary Video 1.** The movie shows the placement of the 5-alanine fragment on the white cartoon of the final structure of crambin. The 2F_o_-F_c_ map of the phases from this initial placement are in blue surfaces at a 1σ contour around the protein with a range of 3 Å.

**Supplementary Video 2.** The movie shows the placement of the 5-alanine fragment on the white cartoon and sticks of the final structure of crambin. The |E| map directly after density modification from this initial fragment in ACORN are in orange surfaces at a 1.5 σ contour around the protein with a range of 3 Å, showing clear, individual spherical densities around individual atoms.

## Notes

### Competing Interest Statement

The authors have declared no competing interest.

## References

1. Wu, M. & Lander, G. C. How low can we go? Structure determination of small biological complexes using single-particle cryo-EM. Current Opinion in Structural Biology 64, 9–16 (2020).

2. Brocchieri, L. & Karlin, S. Protein length in eukaryotic and prokaryotic proteomes. Nucleic acids research 33, 3390–3400 (2005).

3. Nannenga, B. L. & Gonen, T. The cryo-EM method microcrystal electron diffraction (MicroED). Nature Methods 16, 369–379 (2019).

4. Henderson, R. The potential and limitations of neutrons, electrons and X-rays for atomic resolution microscopy of unstained biological molecules. Quart. Rev. Biophys. 28, 171–193 (1995).

5. Rodriguez, J. A. et al. Structure of the toxic core of α-synuclein from invisible crystals. Nature 525, 486–490 (2015).

6. Jones, C. G. et al. The CryoEM Method MicroED as a Powerful Tool for Small Molecule Structure Determination. ACS Cent. Sci. 4, 1587–1592 (2018).

7. Delgadillo, D. A. et al. Microcrystal Electron Diffraction-Guided Discovery of Fungal Metabolites. J. Am. Chem. Soc. (2025) doi:10.1021/jacs.5c01466.

8. Shi, D., Nannenga, B. L., Iadanza, M. G. & Gonen, T. Three-dimensional electron crystallography of protein microcrystals. elife 2, e01345 (2013).

9. Nannenga, B. L., Shi, D., Leslie, A. G. W. & Gonen, T. High-resolution structure determination by continuous-rotation data collection in MicroED. Nature Methods 11, 927–930 (2014).

10. Xu, H. et al. Solving a new R2lox protein structure by microcrystal electron diffraction. Sci. Adv. 5, eaax4621 (2019).

11. Clabbers, M. T. B. et al. MyD88 TIR domain higher-order assembly interactions revealed by microcrystal electron diffraction and serial femtosecond crystallography. Nat Commun 12, 2578 (2021).

12. Nicolas, W. J. et al. Structure of the lens MP20 mediated adhesive junction. Nat Commun 16, 2977 (2025).

13. Jumper, J. et al. Highly accurate protein structure prediction with AlphaFold. Nature 596, 583–589 (2021).

14. Gruene, T. et al. Rapid Structure Determination of Microcrystalline Molecular Compounds Using Electron Diffraction. Angew Chem Int Ed 57, 16313–16317 (2018).

15. Gemmi, M. et al. 3D Electron Diffraction: The Nanocrystallography Revolution. ACS Cent. Sci. 5, 1315–1329 (2019).

16. van Genderen, E. et al. Ab initio structure determination of nanocrystals of organic pharmaceutical compounds by electron diffraction at room temperature using a Timepix quantum area direct electron detector. Acta Cryst A 72, 236–242 (2016).

17. Sawaya, M. R. et al. Ab initio structure determination from prion nanocrystals at atomic resolution by MicroED. Proc. Natl. Acad. Sci. U.S.A. 113, 11232–11236 (2016).

18. Richards, L. S. et al. Fragment-based determination of a proteinase K structure from MicroED data using *ARCIMBOLDO_SHREDDER*. Acta Crystallogr D Struct Biol 76, 703–712 (2020).

19. Martynowycz, M. W., Clabbers, M. T. B., Hattne, J. & Gonen, T. Ab initio phasing macromolecular structures using electron-counted MicroED data. Nature Methods 19, 724–729 (2022).

20. Teeter, M. M. & Hendrickson, W. A. Highly ordered crystals of the plant seed protein crambin. Journal of molecular biology 127, 219–223 (1979).

21. Hendrickson, W. A. & Teeter, M. M. Structure of the hydrophobic protein crambin determined directly from the anomalous scattering of sulphur. Nature 290, 107–113 (1981).

22. Chen, J. C.-H., Hanson, B. L., Fisher, S. Z., Langan, P. & Kovalevsky, A. Y. Direct observation of hydrogen atom dynamics and interactions by ultrahigh resolution neutron protein crystallography. Proceedings of the National Academy of Sciences 109, 15301–15306 (2012).

23. Jelsch, C. et al. Accurate protein crystallography at ultra-high resolution: Valence electron distribution in crambin. Proceedings of the National Academy of Sciences 97, 3171–3176 (2000).

24. Schmidt, A., Teeter, M., Weckert, E. & Lamzin, V. S. Crystal structure of small protein crambin at 0.48 Å resolution. Structural Biology and Crystallization Communications 67, 424–428 (2011).

25. Application note PX031 - Method: MicroED/3D ED on Proteins Using the XtaLAB Synergy-ED. https://rigaku.com/products/crystallography/electron-diffraction/application-notes/3ded031-microed-on-proteins-xtalab-synergy-ed.

26. Usón, I. & Sheldrick, G. M. Advances in direct methods for protein crystallography. Current Opinion in Structural Biology 9, 643–648 (1999).

27. Wennmacher, J. T. C. et al. 3D-structured supports create complete data sets for electron crystallography. Nat Commun 10, 3316 (2019).

28. Martynowycz, M. W., Clabbers, M. T. B., Unge, J., Hattne, J. & Gonen, T. Benchmarking the ideal sample thickness in cryo-EM. Proc. Natl. Acad. Sci. U.S.A. 118, e2108884118 (2021).

29. Tickle, I. J. et al. STARANISO (Global Phasing Ltd., 2018).

30. Xu, H. et al. A Rare Lysozyme Crystal Form Solved Using Highly Redundant Multiple Electron Diffraction Datasets from Micron-Sized Crystals. Structure 26, 667–675.e3 (2018).

31. Hattne, J. et al. Analysis of Global and Site-Specific Radiation Damage in Cryo-EM. Structure 26, 759–766.e4 (2018).

32. Jenkins, H. T. *Fragon* : rapid high-resolution structure determination from ideal protein fragments. Acta Crystallogr D Struct Biol 74, 205–214 (2018).

33. McCoy, A. J. et al. Phaser crystallographic software. Applied Crystallography 40, 658–674 (2007).

34. Dodson, E. J. & Woolfson, M. M. *ACORN* 2: new developments of the *ACORN* concept. Acta Crystallogr D Biol Crystallogr 65, 881–891 (2009).

35. Rodríguez, D. D. et al. Crystallographic ab initio protein structure solution below atomic resolution. Nature Methods 6, 651–653 (2009).

36. Thorn, A. & Sheldrick, G. M. Extending molecular-replacement solutions with SHELXE. Biological Crystallography 69, 2251–2256 (2013).

37. Cowtan, K. The *Buccaneer* software for automated model building. 1. Tracing protein chains. Acta Crystallogr D Biol Crystallogr 62, 1002–1011 (2006).

38. Teeter, M. M., Roe, S. M. & Heo, N. H. Atomic Resolution (0·83 Å) Crystal Structure of the Hydrophobic Protein Crambin at 130 K. Journal of Molecular Biology 230, 292–311 (1993).

39. Weeks, C. M. et al. Crambin: a direct solution for a 400-atom structure. Biological Crystallography 51, 33–38 (1995).

40. Bang, D., Chopra, N. & Kent, S. B. H. Total Chemical Synthesis of Crambin. J. Am. Chem. Soc. 126, 1377–1383 (2004).

41. Chen, J. C.-H. et al. Solvent organization in the ultrahigh-resolution crystal structure of crambin at room temperature. IUCrJ 11, 649–663 (2024).

42. Palatinus, L. et al. Hydrogen positions in single nanocrystals revealed by electron diffraction. Science 355, 166–169 (2017).

43. Kulik, M., Chodkiewicz, M. L. & Dominiak, P. M. Theoretical 3D electron diffraction electrostatic potential maps of proteins modeled with a multipolar pseudoatom data bank. Acta Crystallogr D Struct Biol 78, 1010–1020 (2022).

44. Marques, M. A., Purdy, M. D. & Yeager, M. CryoEM maps are full of potential. Current Opinion in Structural Biology 58, 214–223 (2019).

45. Coppens, P. Comparative X-Ray and Neutron Diffraction Study of Bonding Effects in *s*-Triazine. Science 158, 1577–1579 (1967).

46. Clabbers, M. T., Martynowycz, M. W., Hattne, J. & Gonen, T. Hydrogens and hydrogen-bond networks in macromolecular MicroED data. Journal of Structural Biology: X 6, 100078 (2022).

47. Maki-Yonekura, S., Kawakami, K., Takaba, K., Hamaguchi, T. & Yonekura, K. Measurement of charges and chemical bonding in a cryo-EM structure. Commun Chem 6, 98 (2023).

48. Bücker, R. et al. Serial protein crystallography in an electron microscope. Nat Commun 11, 996 (2020).

49. Liebschner, D., Afonine, P. V., Poon, B. K., Moriarty, N. W. & Adams, P. D. Improved joint X-ray and neutron refinement procedure in Phenix. Biological Crystallography 79, 1079–1093 (2023).

50. Yang, T., Xu, H. & Zou, X. Improving data quality for three-dimensional electron diffraction by a post-column energy filter and a new crystal tracking method. Applied Crystallography 55, 1583–1591 (2022).

51. Clabbers, M. T., Hattne, J., Martynowycz, M. W. & Gonen, T. Energy filtering enables macromolecular MicroED data at sub-atomic resolution. Nature communications 16, 2247 (2025).

52. Vlahakis, N. et al. Fast event-based electron counting for small-molecule structure determination by MicroED. Acta Crystallogr C Struct Chem 81, 116–130 (2025).

53. Martynowycz, M. W., Zhao, W., Hattne, J., Jensen, G. J. & Gonen, T. Collection of Continuous Rotation MicroED Data from Ion Beam-Milled Crystals of Any Size. Structure 27, 545–548.e2 (2019).

54. Martynowycz, M. W. et al. A robust approach for MicroED sample preparation of lipidic cubic phase embedded membrane protein crystals. Nat Commun 14, 1086 (2023).

55. Yamano, A. & Teeter, M. M. Correlated disorder of the pure Pro22/Leu25 form of crambin at 150 K refined to 1.05-A resolution. Journal of Biological Chemistry 269, 13956–13965 (1994).

56. VanEtten, C. H., Nielsen, H. C. & Peters, J. E. A crystalline polypeptide from the seed of Crambe abyssinica. Phytochemistry 4, 467–473 (1965).

57. De La Cruz, M. J., Martynowycz, M. W., Hattne, J. & Gonen, T. MicroED data collection with SerialEM. Ultramicroscopy 201, 77–80 (2019).

58. Mastronarde, D. N. Advanced data acquisition from electron microscopes with SerialEM. Microscopy and Microanalysis 24, 864–865 (2018).

59. CrysAlisPro Software system, version 1.171.38.46. Rigaku Oxford Diffraction (2017).

60. Zheng, S. Q. et al. MotionCor2: anisotropic correction of beam-induced motion for improved cryo-electron microscopy. Nature methods 14, 331–332 (2017).

61. Knudsen, E. B., Sørensen, H. O., Wright, J. P., Goret, G. & Kieffer, J. FabIO: easy access to two-dimensional X-ray detector images in Python. Applied Crystallography 46, 537–539 (2013).

62. Harris, C. R. et al. Array programming with NumPy. Nature 585, 357–362 (2020).

63. Virtanen, P. et al. SciPy 1.0: fundamental algorithms for scientific computing in Python. Nature methods 17, 261–272 (2020).

64. Kabsch, W. XDS. Acta Crystallogr D Biol Crystallogr 66, 125–132 (2010).

65. Kabsch, W. Integration, scaling, space-group assignment and post-refinement. Biological crystallography 66, 133–144 (2010).

66. Winn, M. D. et al. Overview of the *CCP* 4 suite and current developments. Acta Crystallogr D Biol Crystallogr 67, 235–242 (2011).

67. Murshudov, G. N. et al. REFMAC5 for the refinement of macromolecular crystal structures. Biological crystallography 67, 355–367 (2011).

68. Kovalevskiy, O., Nicholls, R. A., Long, F., Carlon, A. & Murshudov, G. N. Overview of refinement procedures within REFMAC5: utilizing data from different sources. Biological Crystallography 74, 215–227 (2018).

69. Grainger, C. T. Pseudo-merohedral twinning. The treatment of overlapped data. Foundations of Crystallography 25, 427–434 (1969).

70. Chen, V. B. et al. MolProbity: all-atom structure validation for macromolecular crystallography. Biological crystallography 66, 12–21 (2010).

71. Schindelin, J., et al. Fiji: an open-source platform for biological-image analysis. Nature methods 9, 676–682 (2012).

72. Meng, E. C. et al. UCSF CHIMERAX : Tools for structure building and analysis. Protein Science 32, e4792 (2023).

73. DeLano, W. L. Pymol: An open-source molecular graphics tool. CCP4 Newsl. Protein Crystallogr 40, 82–92 (2002).

